# High Affinity Nanobodies Block SARS-CoV-2 Spike Receptor Binding Domain Interaction with Human Angiotensin Converting Enzyme

**DOI:** 10.1101/2020.07.24.219857

**Authors:** Thomas J. Esparza, Negin P. Martin, George P. Anderson, Ellen R. Goldman, David L. Brody

**Author notes:** Disclaimer: The views expressed in this manuscript reflect those of the authors, and not those of the Uniformed Services University of the Health Sciences or the Department of Defense.

## Abstract

There are currently no approved effective treatments for SARS-CoV-2, the virus responsible for the COVID-19 pandemic. Nanobodies are 12-15 kDa single-domain antibody fragments that are amenable to inexpensive large-scale production and can be delivered by inhalation. We have isolated nanobodies that bind to the SARS-CoV-2 spike protein receptor binding domain and block spike protein interaction with the angiotensin converting enzyme 2 (ACE2) with 1-5 nM affinity. The lead nanobody candidate, NIH-CoVnb-112, blocks SARS-CoV-2 spike pseudotyped lentivirus infection of HEK293 cells expressing human ACE2 with an EC_50_ of 0.3 micrograms/mL. NIH-CoVnb-112 retains structural integrity and potency after nebulization. Furthermore, NIH-CoVnb-112 blocks interaction between ACE2 and several high affinity variant forms of the spike protein. These nanobodies and their derivatives have therapeutic, preventative, and diagnostic potential.

## INTRODUCTION

Coronaviruses are positive sense, single-stranded RNA viruses. There are seven types of coronaviruses known to infect humans, including the recent 2019 severe acute respiratory syndrome coronavirus 2 (SARS-CoV-2).[1, 2] Patients infected with these viruses develop respiratory symptoms of various severity and outcomes. Since the beginning of the century, there have been three major world-wide health crises caused by coronaviruses: the 2003 SARS-CoV-1 outbreak, the 2012 MERS-CoV outbreak, and the 2019 SARS-CoV-2 outbreak.[3] To date, hundreds of thousands of people have succumbed to the virus during these outbreaks.

The SARS-CoV-2 virus gains entry to human cells via the angiotensin converting enzyme 2 (ACE2) receptor by the interaction of the receptor binding domain (RBD) of the spike protein on the viral surface.[4-8] This RBD-ACE2 interaction provides a clear therapeutic target for binding and prevention of infection (**Fig. 1a**).

**Fig. 1:**
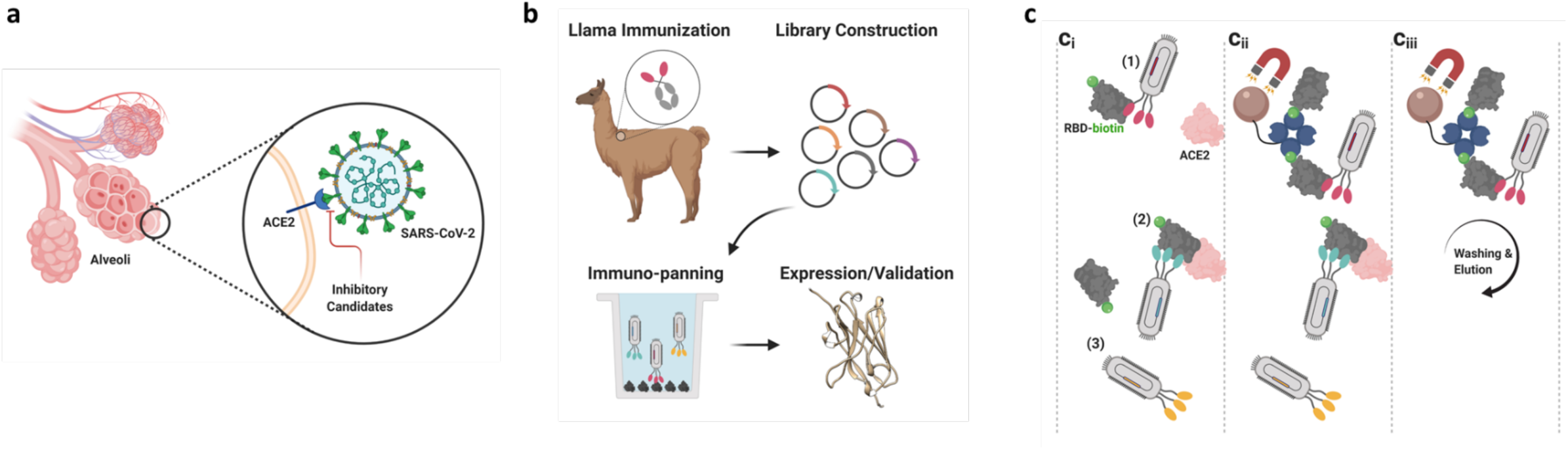
Overview of the therapeutic potential and isolation of nanobodies targeting the SARS-CoV-2 RBD:ACE2 interaction. **a** Illustration of the structure of SARS-CoV-2 spike protein, with receptor binding domain in contact with the human ACE2 receptor on the surface of a lung epithelial cell. A major therapeutic goal is to develop inhibitory agents that disrupt the interaction between the spike protein and the human ACE2 receptor. **b** Isolation of nanobodies binding SARS-CoV-2 spike protein. An adult llama was immunized 5 times over 28 days with purified, recombinant SARS-CoV-2 spike protein. On day 35 after first immunization (7 days after last immunization), llama blood was obtained through a central line, B-cells were isolated, the single heavy-chain variable domains (nanobodies) of the llama antibodies were amplified and cloned to construct a recombinant DNA library containing more than 10^8^ clones. The library of clones was expressed in a phage display format, in which each phage expresses between 1-5 nanobody copies on its surface and contains the DNA sequence encoding that nanobody. Immuno-panning was performed to isolate candidate nanobodies for expression and validation studies. **c** Selection strategy for isolation of nanobody candidates which bind to the RBD:ACE2 interaction surface. Using a phage display library from an adult llama immunized with full length S1 spike protein, nanobodies were isolated which block the interaction between RBD and ACE2. (**c**_**i**_) In a standard radioimmunoassay tube, ACE2 was immobilized and the surface blocked with non-specific protein. Biotinylated-RBD, was incubated with the nanobody phage library and then added to the immuno-tube and allowed to interact with the immobilized ACE2. Biotinylated-RBD with no associated nanobodies, or with nanobody associations which do not block the ACE2 binding domain, bound the immobilized ACE2. (**c**_**ii**_) Biotinylated-RBD with associated nanobodies that blocked the ACE2 binding domain remained in solution and were recovered using streptavidin-coated magnetic particles that bind to the biotin. (**c**_**iii**_) Nanobodies which did not bind to RBD were removed during washing of the magnetic beads. This method allowed for specific enrichment of nanobodies which both bind to the RBD and compete for the RBD-ACE2 binding surface. (Figure generated using BioRender.com)

Many of the antibodies considered for diagnostic or therapeutic applications have been conventional immunoglobulins (IgG). The use of IgGs as therapeutics, while successful in many diseases, is known to have potential pitfalls due to the risk of receptor-mediated immunological reactions.[9] Treatment or prophylaxis of a pulmonary virus can be delivered via aerosolization and inhalation; thus, the size and biophysical characteristics are paramount considerations.

The camelid family, which includes llamas, produce additional subclasses of IgGs which possess an unpaired heavy-chain variable domain.[10-12] This heavy-chain variable domain has demonstrated the ability to function as an independent antigen-binding domain with similar affinity as a conventional IgG. These heavy chain variable domains can be expressed as a single domain, known as a VHH or nanobody, with a molecular weight 10% of the full IgG. Nanobodies generally display superior solubility, solution stability, temperature stability, strong penetration into tissues, are easily manipulated with recombinant molecular biology methods, and possess robust environmental resilience to conditions detrimental to conventional IgGs.[13-15] In addition, nanobodies are weakly immunogenic which reduces the likelihood of adverse effects compared to other single domain antibody such as those derived from sharks or synthetic platforms. Successful framework modifications have been implemented to humanize VHH sequences without significant alteration of biophysical properties.[16] Importantly, the recent approval of the first-in-class nanobody treatment for thrombotic thrombocytopenic pupura, Caplacizumab, bolsters the therapeutic potential of VHH derivatives.[17] Therefore, nanobodies that bind to the SARS-CoV-2 RBD and block the ACE2 interaction are an attractive therapeutic for prevention and treatment of viral infection.[18]

## RESULTS

Using standard methods to immunize llama, B-cell nanobody DNA sequences were isolated and a phage display library with over 10^8^ clones was created (**Fig. 1b**). From this phage library, 13 unique lead candidate nanobodies that bind to the SARS-CoV-2 spike protein RBD were isolated, several of which block the spike protein-ACE2 interaction with high potency.

To isolate the candidate nanobodies, a novel screening strategy was designed and executed to specifically select for nanobodies that not only bound to the SARS-CoV-2 spike RBD, but also interfered with the interaction with the human ACE2 protein. The purity of *in vitro* binding of commercially available recombinant spike protein RBD and human ACE2 protein (**Supplementary Fig. 1a-c**) that were used for the screening strategy was validated. In this screening strategy (**Fig. 1c**), recombinant human ACE2 protein was immobilized in tubes. Then, biotinylated-RBD was incubated with the nanobody phage library and allowed to interact with the immobilized ACE2. Biotinylated-RBD with no associated nanobodies, or with nanobody associations that did not block the ACE2 binding domain, bound the immobilized ACE2. Biotinylated-RBD with associated nanobodies that blocked the ACE2 binding domain remained in solution and were recovered using streptavidin-coated magnetic particles that bind to the biotin. Nanobodies that did not bind to RBD were removed during washing of the magnetic beads. This method allowed for specific enrichment of nanobodies that bind to the RBD and compete for the RBD-ACE2 binding surface.

Using this novel screening strategy, hundreds of phage clones were isolated, which when sequenced, reveal 13 unique full length nanobody DNA sequences, termed NIH-CoVnb-101 through NIH-CoVnb-113 (**Fig. 2**). These sequences were distinct from the previously published sequences that also bind SARS-CoV-2 spike protein.[19-26] Additional reports demonstrate compelling binding to SARS-CoV-2 spike protein, yet have not disclosed nanobody sequences allowing for a direct comparison.[27-30] The complementarity determining region (CDR) 3 domain responsible for much of the specific binding of nanobodies to their targets, in NIH-CoVnb-112 and the other new nanobodies were generally longer than those currently reported. These findings indicated that novel nanobody DNA sequences were isolated.

**Fig. 2:**
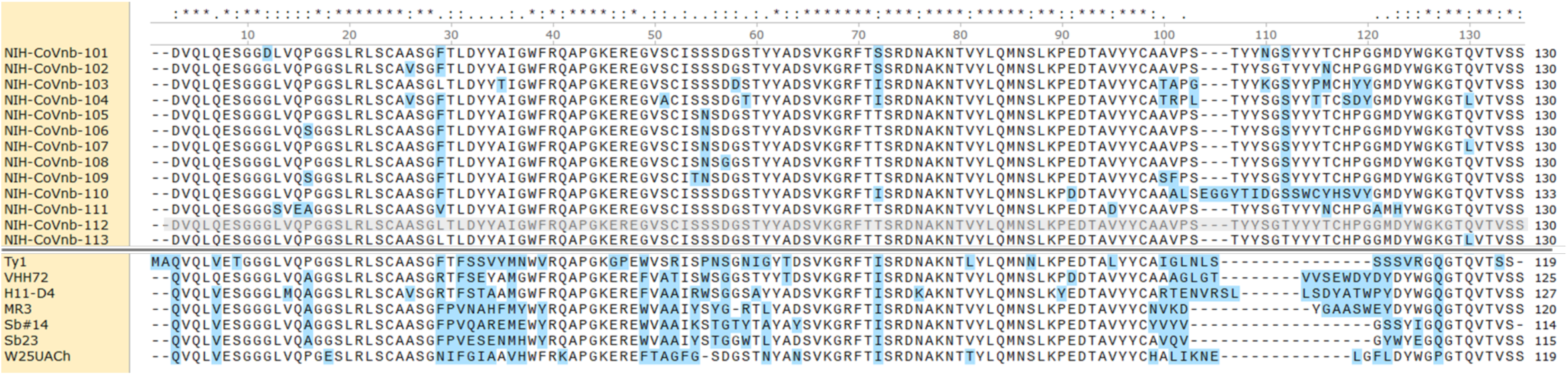
Protein sequences for novel nanobodies that bind to the SARS-CoV-2 spike protein receptor binding domain. Single letter amino acid codes. Clustal Omega algorithm used for alignment. Blue highlights indicate sequence diversity with NIH-CoVnb-112, highlighted in gray, set as the reference sequence for comparison. For comparison, seven previously reported nanobody sequences have clearly distinct sequences: Ty1[23], VHH72[24], H11-D4[19], MR3[20], Sb#14[21], Sb23[22], and W25UACh[25] and possess shorter CDR3 domains (represented in NIH-CoVnb-112 by amino acids 99-120).

Representatives from each of these unique nanobody sequences were produced in bacteria, purified, and tested for binding to SARS-CoV-2 spike RBD (**Fig. 3**). All of the nanobodies bound to recombinant SARS-CoV-2 spike RBD with high affinity as measured by Octet biolayer interferometry. The strongest binding nanobody was NIH-CoVnb-112 (**Fig. 3e**) with an affinity of 4.9 nM. NIH-CoVnb-112 had both a fast on rate (1.3e5/M/sec.) and a slow off rate (6.54e-4/sec.), with kinetics compatible with 1:1 binding. In a complementary measurement, ProteOn XPR36 surface plasmon resonance provided good agreement (**Supplementary Fig. 2**) with an affinity of 2.1nM. These results indicate that the novel nanobodies bind to the SARS-CoV-2 spike RBD with very high affinity. The long CDR3 in NIH-CoVnb-112 and the other new nanobodies may in part underlie their extraordinarily high affinity, though this is clearly not the only factor in that the nanobody with the longest CDR3, NIH-CoVnb-110, is not one of the top 5 highest affinity nanobodies. A portion of the high affinity may be explained by the presence of a non-canonical disulfide bond between the CDR3 and the framework, which may provide additional stability.[10] Measurements using circular dichroism (CD) during heating (**Supplementary Fig. 3a**) reveal that the nanobody structure resists unfolding until 74.4°C (**Supplementary Fig. 3b**) and upon cooling 73% of the structure returned to the baseline CD value. These data support an extremely stable, robust, high affinity nanobody.

**Fig. 3:**
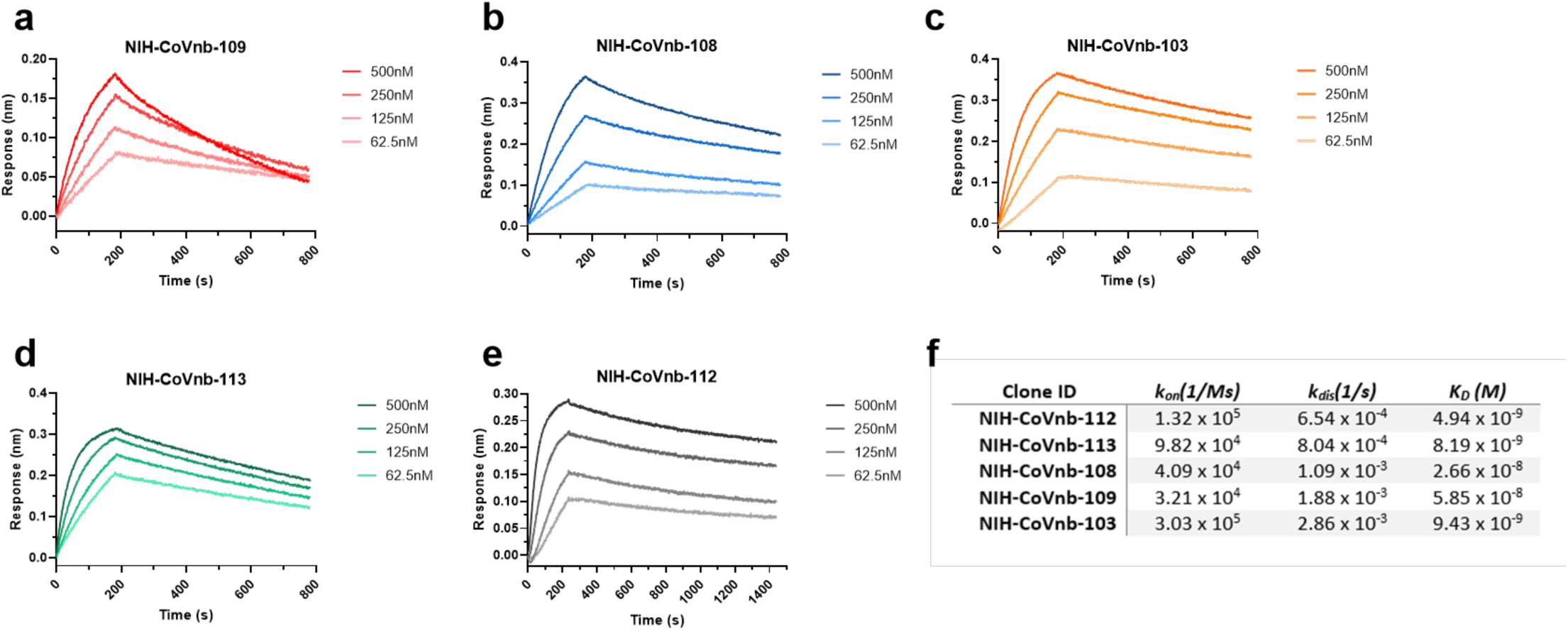
Affinity binding curves of isolated anti-SARS-CoV-2 RBD nanobodies. Using Biolayer Interferometry on a BioForte Octet Red96 system, association and dissociation rates were determined by immobilizing biotinylated-RBD onto streptavidin coated optical sensors (**a-e**). The RBD-bound sensors were incubated with specific concentrations of purified candidate nanobodies for a set time interval to allow association. The sensors were then moved to nanobody-free solution and allowed to dissociate over a time interval. Curve fitting using a 1:1 interaction model allows for the affinity constant (K_D_) to be measured for each nanobody as detailed in (**f**).

In an important validation of the *in vitro* efficacy of the candidate nanobodies, the nanobodies were able to interfere with SARS-CoV-2 spike RBD binding to human ACE2 protein (**Fig. 4a**). Specifically, a competitive inhibition assay was designed and implemented, in which recombinant RBD was coated onto Enzyme Linked Immuno-Sorbent Assay (ELISA) plates and soluble ACE2 binding was assessed. Without any interference, soluble ACE2 binding was indicated by high colorimetric absorbance. At increasing concentrations, each of the new nanobodies, showed progressively less ACE2 binding. For the most potent anti-SARS-CoV-2 RBD nanobody, NIH-CoVnb-112, the concentration at which 50% of the ACE2 binding was blocked (half maximal effective concentration; termed EC_50_) was found to be 0.02 micrograms/mL, equivalent to 1.11 nM. The rank order of ACE2 competition EC_50_ matched the rank order of RBD binding affinities for the novel nanobodies assessed. As an additional confirmation, the experiment was repeated using a commercially available SARS-CoV-2 spike RBD- human ACE2 protein competitive inhibition assay (GenScript). The results were very similar to those obtained using the initial assay (**Fig. 4b**). These results indicate that the novel nanobodies block SARS-CoV-2 spike protein RBD binding to ACE2, an essential human receptor responsible for viral infection.

**Fig. 4:**
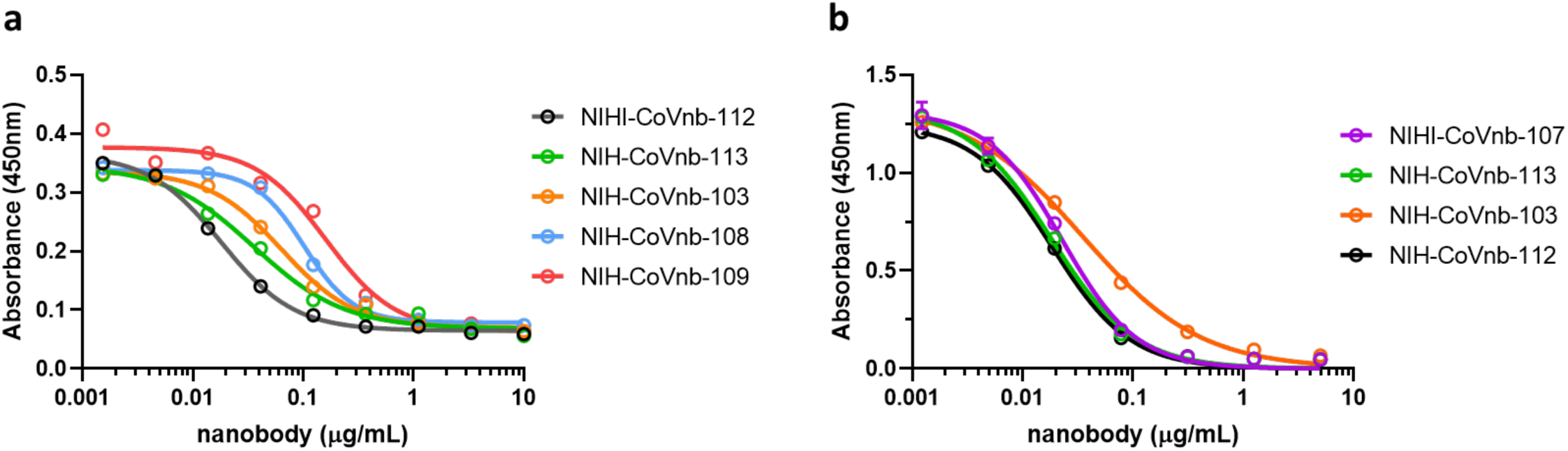
Competitive inhibition of ACE2 and RBD binding using anti-SARS-CoV-2 RBD nanobodies. **a** RBD coated ELISA plates were blocked with non-specific protein and incubated with serial dilutions of each candidate anti-SARS-CoV-2 RBD nanobody. Biotinylated-ACE2 was added to each well and allowed to bind to unoccupied RBD. The ELISA was then developed using a standard streptavidin-HRP and tetramethylbenzidine reaction. Unoccupied RBD allows for a positive reaction signal which is suppressed in the presence of bound competitive nanobody. NIH-CoVnb-112 produces the greatest inhibition of ACE2 binding with an EC_50_ of 0.02 micrograms/mL (1.11 nM). **b** Comparable findings using the commercially available Genscript SARS-CoV-2 neutralization surrogate assay.

There have been many variants of the spike protein RBD described recently that increase the binding affinity to the human ACE2 receptor.[31] Several of these variants including N354D D364Y, V367F, and W436R have been reported to have up to 100 fold higher affinity for ACE2 than the prototype RBD *in vitro*. NIH-CoVnb-112 blocked interaction between human ACE2 and three of these variants with similar EC_50_ compared to its blocking effects on the prototype sequence spike protein RBD (**Fig. 5b**). Binding affinity of NIH-CoVnb-112 to the variants was also similar to that of the prototype sequence (**Fig. 5a**). Thus, NIH-CoVnb-112 may provide robust blocking of SARS-CoV-2 spike protein RBD binding to ACE2.

**Fig. 5:**
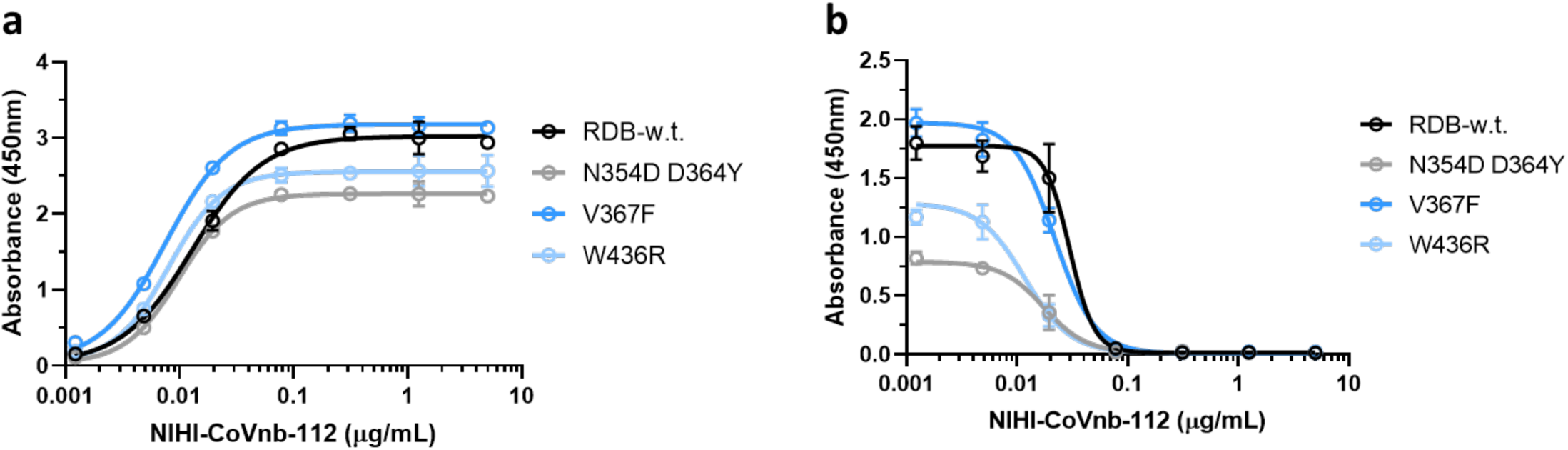
Interaction of NIH-CoVnb-112 with SARS-Cov-2 Spike Protein RBD variants. **a** Binding of NIH-CoVnb-112 to RBD “wild type” and 3 variant forms of the RBD had similar affinity, with half maximal binding at approximately 0.01 micrograms/mL **b** Competitive inhibition assay: RBD “wild type” and 3 variant forms of the RBD coated ELISA plates were blocked with non-specific protein and incubated with dilutions of each the lead candidate anti-SARS-CoV-2 RBD nanobody NIH-CoVnb-112. Biotinylated-ACE2 was added to each well and allowed to bind to unoccupied RBD. The ELISA was then developed using a standard streptavidin-HRP and tetramethylbenzidine reaction. Unoccupied RBD allows for a positive reaction signal which is suppressed in the presence of bound competitive nanobody. NIH-CoVnb-112 inhibited ACE2 binding to each of the variants with a similar EC_50_ of 0.02 micrograms/mL (1.11 nM).

We then assessed whether NIH-CoVnb-112 binds to the same spike protein RBD epitope as the previously reported nanobody VHH72 [24]. We reasoned that if both nanobodies bind to the same epitope, their signals would occlude each other when applied at saturating concentrations on bilayer interferometry. In contrast, if they bind to different epitopes, their signals would be additive on bilayer interferometry. We found that when NIH-CoVnb-112 was applied at 500 nM (>100 x greater than the K_D_) for long enough to reach steady state, and then VHH72 was applied, the signals were clearly additive (**Supplementary Fig. 5a**). This result clearly indicates that NIH-CoVnb-112 binds to a SARS-CoV-2 spike protein RBD epitope that is distinct from that of VHH72. This result is consistent with the reported findings that VHH72 recognizes a non-ACE2 binding motif on the spike protein [24], whereas NIH-CoVnb-112 has RBD-ACE2 interaction disruption potency that is quantitatively similar to its affinity- a result that is most consistent with NIH-CoVnb-112 binding directly to the ACE2 interaction domain.

We then performed additional tests to determine if NIH-CoVnb-112 possesses the ability to also bind to the SARS-CoV-1 spike protein RBD domain. We prepared a biotinylated version of NIH-CoVnb-112 which could be used in parallel to test binding to several targets. Biotinylated NIH-CoVnb-112 was immobilized onto streptavidin Octet sensors and then allowed to associate (**Supplementary Fig. 5b**) with either SARS-CoV-1 RBD or SARS-CoV-2 RBD at 500 – 125nM concentrations. As previously demonstrated, NIH-CoVnb-112 binds robustly to SARS-CoV-2 RBD, yet we found that there was essentially no binding to SARS-CoV-1 RBD. In an orthogonal confirmation, we performed an ELISA in which each RBD was coated onto a standard ELISA plate and then allowed to incubate with NIH-CoVnb-112. The ELISA results were similar to the Octet results: (**Supplementary Fig. 5c**) NIH-CoVnb-112 binds strongly to SARS-CoV-2 RBD and does not associate with SARS-CoV-1 RBD. These results clearly indicate that NIH-CoVnb-112 does not cross react and bind to SARS-CoV-1 spike protein RBD.

The presence of bacterial endotoxins introduces an unwanted liability during the production of NIH-CoVnb-112 due to expression in *E*. *coli*. To remove this risk, we cloned the coding sequence for the nanobody into an expression vector for the methylotrophic yeast *Pichia pastoris*. The expression vector, pICZalpha, contains an alpha-factor secretion signal sequence which directs the expressed protein into the media supernatant. Following selection of a positive recombinant clone we found an expression yield of 40 milligrams per liter following 96 hours of methanol batch-fed culture.

We envision therapeutic utility for NIH-CoVnb-112 as a nebulized deliverable treatment to the respiratory tract of patients. Owing to the small size and general stability of the nanobody, we sought to test if we could successfully nebulize, recover, and confirm the stability of NIH-CoVnb-112 relative to non-nebulized nanobody. Using the commercially available Aerogen® Solo vibrating mesh nebulizer, we nebulized Pichia expressed NIH-CoVnb-112 into an in line custom bead condenser (**Fig. 6a**) to collect the post-nebulized nanobody. We recovered greater than 90% of the total input protein mass. Following nebulization recovery, we incubated the pre-nebulization and post-nebulization samples at 37°C for 24hr to provide physiologic temperature exposure. When assessed by SDS-PAGE gel (**Fig. 6b**) we found that the pre-nebulization and post-nebulization NIH-CoVnb-112 appear identical with no evidence of degradation or aggregation products. Similarly, analysis by size-exclusion chromatography (**Fig. 6c**) on a Superdex 75 column yielded similar elution profiles with no evidence of degradation or aggregation relative to the pre-nebulization sample. We conclude that NIH-CoVnb-112 is resilient to degradation or aggregation during nebulization.

**Fig. 6:**
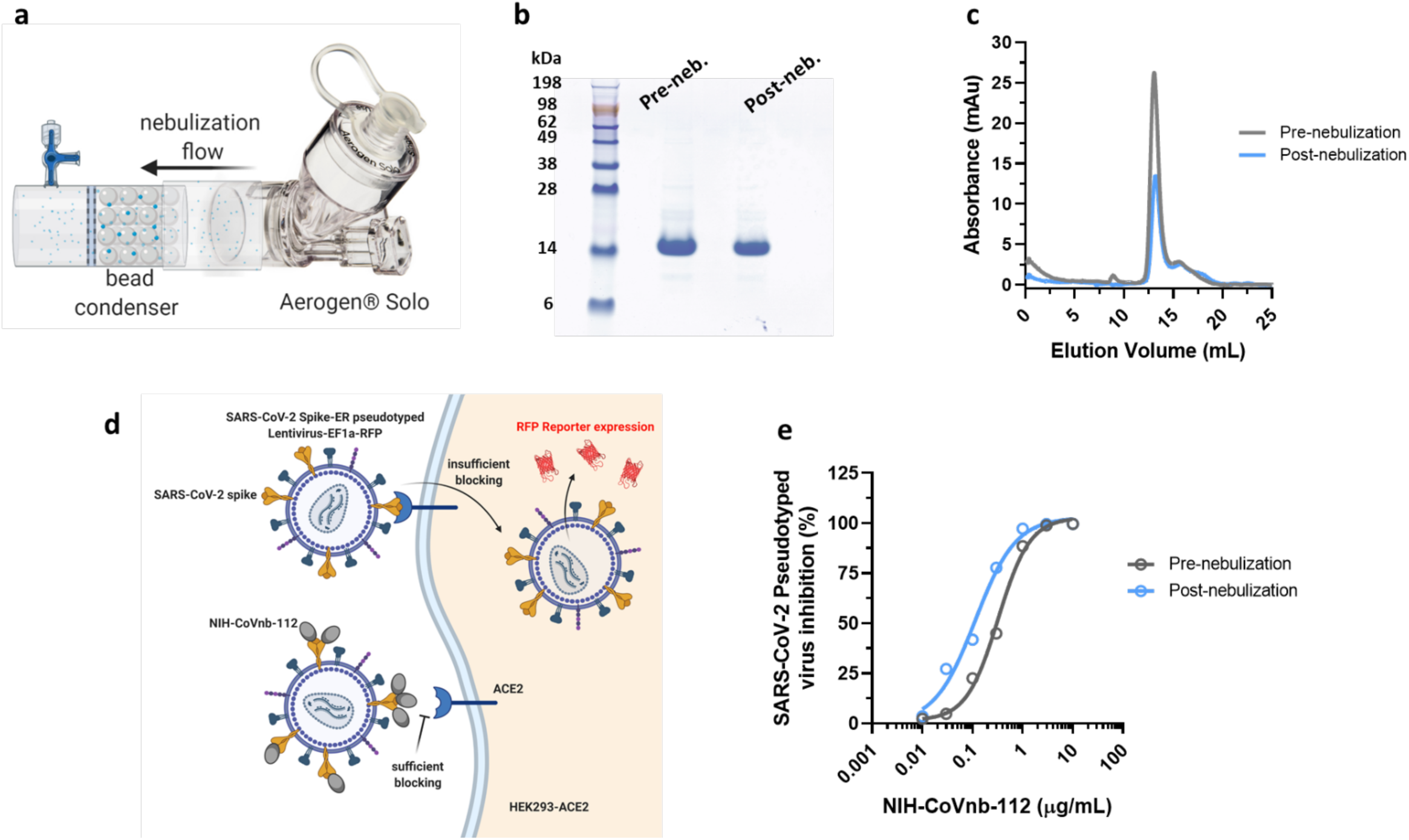
NIH-CoVnb-112 stability and potent inhibition of SARS-CoV-2 pseudovirus following nebulization. **a** An Aerogen® Solo High-Performance Vibrating Mesh nebulizer was placed in line with a custom glass bead condenser to allow for collection of the nanobody following nebulization. **b** A pre-nebulization and post-nebulization sample of NIH-CoVnb-112 was analyzed by SDS-PAGE gel. The dominant band for each sample remains at approximately 14kDa indicating no detectable degradation or aggregation of NIH-CoVnb-112 following nebulization. **c** Size exclusion chromatography demonstrated a prominent peak eluting at 13.5mL elution volume in both pre and post-nebulization samples. The other minor profile peaks are evident in both samples, with an absence of unique peaks which would indicate a change in conformation state or degradation. **d** A fluorescence reporter assay utilizing SARS-CoV-2 spike protein pseudotyped lentivirus was used to demonstrate potent inhibition of the RBD:ACE2 interaction. HEK293 cells overexpressing human ACE2 were cultured for 24 hours with pseudotyped virus which was pretreated with NIH-CoVnb-112 at different concentrations. Inhibition of the spike RBD occurs when the virus is not able to transduce the HEK293-ACE2 cells and subsequently produce RFP reporter protein. **e** Following 48hr incubation, HEK293-ACE2 cells were analyzed by flow cytometry to quantify the fluorescence level. NIH-CoVnb-112 potently inhibited viral transduction both pre and post-nebulization with an EC_50_ of 0.323 micrograms/mL (23.1nM) and 0.116 micrograms/mL (8.3nM) respectively. (Figure elements generated using BioRender.com)

To further assess the stability of NIH-CoVnb-112, we performed incubation in pooled normal human plasma and recombinant human albumin followed by affinity measurement to assess preservation of SARS-CoV-2 RBD binding potential. NIH-CoVnb-112, at 5µM, was incubated for 24 or 48hrs in the presence of human plasma, which contains proteases, or recombinant human albumin alone at 37°C with gentle mixing. In addition, for each condition a time zero control was prepared to account for potential matrix effects. The samples were diluted to 500nM based on the NIH-CoVnb-112 starting concentration and assessed by biolayer interferometry for binding to SARS-CoV-2 RBD. NIH-CoVnb-112 binding (**Supplementary Fig. 4a**) SARS-CoV-2 RBD following treatment in pooled human plasma at time zero, 24hr, and 48hr had negligible impact on binding. Similarly, NIH-CoVnb-112 binding (**Supplementary Fig. 4b**) SARS-CoV-2 RBD following treatment in recombinant human albumin at all time points had no apparent effect on binding. While further *in vivo* characterization is necessary, these data support the interpretation that NIH-CoVnb-112 is satisfactorily stable in the presence of plasma.

Finally, while in a plate-based protein inhibition assay NIH-CoVnb-112 robustly inhibits the SARS-CoV-2 RBD and ACE2 interaction, we sought to demonstrate if a similar inhibition is achieved in a viral transduction assay. We used pseudotyped SARS-CoV-2 spike protein bearing lentivirus with an RFP reporter system to test if NIH-CoVnb-112 could inhibit the transduction of human embryonic kidney cells (HEK293T/17) overexpressing human ACE2 (HEK293-ACE2). Transduction of HEK293-ACE2 cells results in RFP expression (**Fig. 6d**) in the absence of an inhibitory agent. The pseudotyped virus were produced using a human codon optimized SARS-CoV-2 spike protein sequence with the endoplasmic reticulum retention signal removed. Pseudotyped virus were purified over a sucrose cushion and viral titer determined using digital droplet PCR. A serial dilution of Pichia expressed, pre-nebulization and post-nebulization NIH-CoVnb-112 samples were incubated with a Multiplicity of Infection (MOI) of 0.5 followed by incubation on HEK293-ACE2 cells for 48hrs. Following transduction, brightfield and epifluorescence images were acquired for both pre-nebulization (**Supplementary Fig. 6a**) and post-nebulization (**Supplementary Fig. 6b**) conditions. Incubation at or above 0.3 micrograms/mL NIH-CoVnb-112 results in robust inhibition for both conditions. Following trypsinization and fixation, the individual conditions were analyzed by flow cytometry to determine the percentage of RFP expressing cells. The population of single events were recorded for both pre-nebulization (**Supplementary Fig. 7a**) and post-nebulization (**Supplementary Fig. 7b**) conditions. The number of RFP positive events for each condition were normalized relative to the virus-alone control to yield a percent inhibition of the SARS-CoV-2 pseudotyped lentivirus (**Fig. 6e**). The pre-nebulization NIH-CoVnb-112 has an EC_50_ of 0.323 micrograms/mL (23.1nM) while the post-nebulization NIH-CoVnb-112 has an EC_50_ of 0.116 micrograms/mL (8.3nM). The difference in EC_50_ values between the two conditions is most likely an effect of assay variance and we do not speculate that nebulization produced an increase in potency. Thus, we find that NIH-CoVnb-112 potently inhibits viral transduction in an infection relevant pseudotyped SARS-CoV-2 virus model.

## DISCUSSION

Here, we report several nanobodies that bind to the SARS-CoV-2 spike protein RBD and block spike protein interaction with the ACE2 receptor. The affinity of NIH-CoVnb-112 in monomeric form is substantially better than the monomeric form of previously reported candidate nanobody therapeutics for SARS-CoV-2: 39 nM for monomeric VHH72[24], ∼50 nM for monomeric Ty1[23], 30.7nM for monomeric Sb#14[21], 10nM for monomeric Sb23[22], and incompletely reported but likely lower than 1B and 3F[27]. The affinity and blocking potency of NIH-CoVnb-112 will likely be even higher when formulated into dimeric, trimeric, or other protein engineered constructs.[27, 30, 32, 33]

Importantly, these nanobody therapeutics may be delivered via inhalation.[34] Inhalation has major advantages over other routes of administration and could be one of the most important potential uses for a nanobody therapeutic for SARS-CoV-2. The most directly analogous study was performed by Larios Mora et al., who showed that nebulized treatment of newborn lambs with ALX-0171 reduced clinical, virological, and pathological manifestations of respiratory syncytial virus.[35] ALX-0171 is a trimeric nanobody therapy produced in *Pischia pastoris* that binds to the respiratory syncytial virus fusion protein. Nebulization was performed using a commercially available human-use Aeroneb Solo system (https://www.aerogen.com/aerogen-solo-3/), interfaced with a nose mask. The Aeroneb Solo nebulizer produced ∼3.3 micron particles, appropriate for deep lung delivery. The authors demonstrated that once daily nebulization resulted in concentrations of ALX-0171 in lung epithelial lining fluid that were >10 times higher than the *in vitro* EC_50_ for respiratory syncytial virus. All treated lambs had undetectable infectious virus in lung epithelial lining fluid, which supports that the proposed route of administration could reduce infectivity. After nebulization, blood concentrations were ∼1,000 fold lower than lung epithelial lining fluid concentrations, which is important because low blood concentrations likely reduce systemic toxicity and risk of host antibody formation. Of note, the parent monomeric nanobody to the respiratory syncytial virus fusion protein had an affinity of 17 nM, whereas the nanobody trimer ALX-0171 had an affinity of 0.113 nM. The trimeric ALX-0171 was produced by fermentation in *Pischia pastoris* with a yield of 7.5 grams per liter.[32] Thus, there is great potential for an inhaled therapeutic for SARS-CoV-2 derived from the NIH-CoVnb-112 nanobody, and other similar nanobodies. There are many other potential therapeutic uses for the NIH-CoVnb-112 nanobody. These include intravenous, intramuscular, or subcutaneous treatment formulations for early stage disease, as well as prevention in high risk individuals.

It appears that late-stage disease is mediated more by immunological responses and less by virological pathogenesis, making it important to initiate virologically-targeted therapy early. Importantly, it is likely that nanobody treatments would be substantially less expensive to produce than conventional monoclonal antibodies such as those currently in clinical development pipelines (e.g. REGN-CoV-2 “antibody cocktail” https://www.regeneron.com/covid19 and LY-CoV555: https://investor.lilly.com/news-releases/news-release-details/lilly-begins-worlds-first-study-potential-covid-19-antibody). Thus, it would be reasonable to initiate treatment with an inexpensive and widely available therapeutic early, even before it is clear whether or not an infection will become severe. Furthermore, even if vaccines are successfully developed, preventatives and treatments for acute illness would be beneficial, as vaccines may not be 100% effective, especially in those who do not mount adequate immune responses. By analogy, oseltamivir (Tamiflu) is an essential medicine even in the context of successful influenza vaccines. Thus, there is a substantial need for inexpensive and safe preventative and acute candidate treatments such as the NIH-CoVnb-112 nanobody.

In addition to therapeutic applications, an inexpensive binding reagent such as NIH-CoVnb-112 nanobody could be used diagnostically. Antibody-based tests could be used to assess for SARS-CoV-2 spike protein in body fluids and the environment. The low cost and temperature stability of nanobodies would be advantageous in this setting.

Substantial effort will be required to fully characterize and develop NIH-CoVnb-112. We have demonstrated successful neutralization assays involving pseudotyped lentivirus and authentic SARS-CoV-2 isolate neutralization assays are currently underway. Future activities include performing detailed structural and biophysical characterization involving Cryo-EM, X-ray crystallography, *in vivo* pharmacokinetics and stability assessment, immunogenicity determination, framework humanization, assessing multimerized formats, and further storage and delivery stability assessments. It is encouraging that NIH-CoVnb-112 binds to and blocks three recently reported RBD variants, but it will be of great interest to determine whether the candidate nanobodies provide broad spectrum blocking against other variant forms of the SARS-CoV-2 spike proteins.[31] Additional assays will be required to determine whether NIH-CoVnb-112 protects against mutation-based viral escape alone or as part of a therapeutic cocktail.

Animal testing will be critically important. The optimized nanobody construct will be tested for *in vivo* safety and efficacy in an established small animal model of SARS-CoV-2 infection such as human ACE2 transgenic mouse or Syrian hamster.[36, 37] Subsequently, testing for *in vivo* safety and efficacy in an established large animal model of SARS-CoV-2 infection such as rhesus macaque or baboon will be conducted to further support an investigational new drug application.[38] While other nanobodies have been produced in large scale current Good Manufacturing Practice facilities by fermentation, there is no guarantee that NIH-CoVnb-112 constructs can be produced in sufficient quantities at reasonable cost. Exploration of the manufacturing parameters will be critical for product development.

There are several limitations of the current results. First, the characterization of NIH-CoVnb-112 and the other nanobodies is not yet complete. Under the current circumstances, we have opted to move forward with dissemination of our findings earlier than might otherwise be typical. Second, a potential new therapeutic such as this nanobody would, even under the best circumstances, come to market relatively slowly compared to a repurposed approved drug. Thus, it is possible that if an approved drug is found to be effective against SARS-CoV-2, it will be available long before this or any other nanobody candidate therapy. Third, it is possible that NIH-CoVnb-112 and other nanobodies may provoke host immune responses. NIH-CoVnb-112 is a llama protein fragment that is similar to human proteins yet cannot be fully humanized and maintain desirable biophysical characteristics. Fourth, NIH-CoVnb-112 and other nanobodies may have unexpected toxicity. There is only one United States Food and Drug Administration-approved nanobody therapeutic, Caplacizumab, and there has not been exposure to a sufficient number of human patients to fully characterize the safety of this class of therapeutics. Despite these limitations, the novel nanobodies that bind to the spike protein RBD have potential widespread utility in the battle against the SARS-CoV-2 pandemic.

## MATERIALS AND METHODS

### Immunization of Llamas

An adult male 16-year-old llama, named Cormac, was immunized under contract at Triple J Farms (Kent Laboratories, Bellingham, WA). Subcutaneous injection was performed at multiple locations with 100 µg of SARS-CoV-2 S1 protein (S1N-C5255, ACRO Biosystems) emulsified in complete Freund’s adjuvant on day 0, followed by additional 100 µg immunizations emulsified with incomplete Freund’s adjuvant on days 7, 14, 21, and 28. On day 35, peripheral blood was isolated and shipped on ice for further processing. Triple J Farms operates under established National Institutes of Health Office of Laboratory Animal Welfare Assurance certification number A4335-01 and United States Department of Agriculture registration number 91-R-0054.

### Generation of Immune Single-domain Antibody Phage Display Library

The general synthesis methods utilized are adapted from work by Pardon et al.[39] Peripheral blood mononuclear cells (PBMCs) were isolated from llama whole blood using Uni-Sep Maxi density gradient tubes (#U-10, Novamed) according to the manufacturer’s directions. The platelet-rich plasma was further processed and stored at −80 °C for use in measuring antibody titer. The PBMCs were rinsed with 1x phosphate buffered saline (PBS), aliquoted, and frozen at −80 °C prior to use. For total RNA isolation a minimum of 1×10^7^ PBMCs were processed using a RNeasy Mini Kit (Qiagen). First-strand synthesis of complimentary DNA from the resulting total RNA was prepared using the SuperScript™ IV First-Strand Synthesis kit (#18091050, Invitrogen) with random hexamer primed conditions and 2 µg total RNA. Synthesized first-strand complimentary DNA was used to amplify the heavy-chain variable domains using Q5 high-fidelity DNA polymerase (New England Biolabs) and the described primers (CALL001: 5’-GTCCTGGCTGCTCTTCTACAAGG-3’ and CALL002: 5’-GGTACGTGCTGTTGAACTGTTCC-3’). The resulting amplified variable domains were separated on a 1.2% (w/v) low melting point agarose gel and the approximately 700 base pair band corresponding to the heavy chain only immunoglobulin was extracted and purified using QIAquick Gel Extraction Kit (Qiagen). A secondary amplification of the VHH domain was performed using modified primers (VHH-Esp-For: 5’-CCGGCCATGGCTGATGTGCAGCTGCAGGAGTCTGGRGGAGG-3’ and VHH-Esp-Rev: 5’-GTGCGGCCGCTGAGGAGACGGTGACCTGGGT-3’) based on those used by Pardon et al. to introduce cloning compatible sequence for the phagemid pHEN2.[39] The pHEN2 phagemid allows for in-frame cloning of the VHH sequences with the pIII M13 bacteriophage gene and the inclusion of a C-terminal 6xHis tag and triple myc tag separated from the pIII sequence by an amber stop codon (TAG). The amplified VHH sequences were then restriction endonuclease digested using NcoI and NotI (New England Biolabs) and purified from the reaction components. The resulting digested VHH sequences were ligated into NcoI/NotI digested pHEN2 phagemid at a 3:1 (insert:phagemid) ratio overnight at 16 °C followed by purification and then electroporation into phage-display competent TG-1 cells (#60502-1, Lucigen). The library was plated onto 2xYT agar plates containing 100 µg/mL carbenicillin and 2% glucose at 37 °C overnight. The resulting library contained >10^8^ independent clones. The library was pooled and archived as glycerol stocks. Standard phage amplification of the representative library was performed using M13KO7 helper phage (#18311019, Invitrogen) followed by precipitation with 20% polyethylene glycol 6000 /2.5M sodium chloride on ice to purify the phage particles for downstream immunopanning. All post-reaction purifications utilized the MinElute PCR Purification Kit (Qiagen).

### Competitive Immunopanning

Selection of nanobodies that block the RBD-ACE2 interaction was performed using a bias-selection strategy. Standard radioimmunoassay tubes were coated with 500 µL of human ACE2 protein (#AC2-H52H8, ACRO Biosystems) solution at 5 µg/mL in sodium carbonate buffer, pH 9.6 overnight at 4 °C. The coating solution was removed, and the tube blocked with a 2% (w/v) non-specific blocking solution which was alternated (bovine serum albumin, nonfat dry milk, or Li-Cor Intercept) each panning round to reduce enrichment of off-target clones. Amplified phage library (approximately 10^11^ phage) was mixed 1:1 with the relevant blocking solution to yield a 500 µL volume to which 1µg biotinylated RBD (#SPD-C82E9) was added and allowed to associate for 30 minutes at room temperature with mixing at 600 rpm. The phage library complexed with biotinylated-RBD was then transferred to the blocked radioimmunoassay tube and allowed to bind for 30 minutes at room temperature with mixing at 600 rpm. The non-associated biotinylated-RBD was then recovered by addition of 10µL Dynabeads™ M-270 Streptavidin for 10 minutes and the supernatant transferred to a new blocked-microcentrifuge tube. The magnetic beads were washed 10 times with 1xPBS and the bound phage eluted with 100mM triethylamine solution and neutralized with 1:10 volume 1M Tris-HCl, pH 8.0. The resulting phage elution was used to infect TG-1 cells and additional phage amplification and immunoselections performed.

### Phage Display Clone Screening

Individual colonies were selected from the phage display enrichment plates and cultured in 2xYT-carbenicillin containing microbial media in a 96-deep well block at 37 °C with 300 rpm shaking for 4-6 hours. Expression of nanobody was induced by addition of isopropyl-beta-D-thiogalactoside (GoldBio) to a final concentration of 1mM and continued incubation overnight. The cells were pelleted by centrifugation at 1,000xG for 20 minutes. The media supernatant was removed, and the block placed at −80 °C for 1 hour. The block was then allowed to equilibrate to room temperature for 15 minutes, 500 µL 1xPBS was added, and the block agitated at 1,500 rpm for 1 hour to allow for release of nanobody from the periplasmic space. The block was then centrifuged for 20 minutes at 2,000xG. Nunc Maxisorp plates were coated with SARS-CoV-2 RBD (SPD-C52H3, ACRO Biosytems) as described above and blocked with 2% bovine serum albumin (BSA). Nanobody containing supernatant was transferred to the blocked plate and incubated for 1 hour at room temperature. The assay plate was washed and 100 µL peroxidase conjugated goat anti-alpaca VHH domain specific antibody (#128-035-232, Jackson ImmunoResearch) transferred into each well and incubated for 1 hour at room temperature. After a final wash the plate was developed by addition of 100 µL tetramethylbenzidine (TMB) (#T5569, Sigma-Aldrich). Assay development was stopped by addition of 50µL 1M sulfuric acid and the assay absorbance measured at 450 nm on a Biotek Synergy 2 plate reader. The assay plate was washed with 1xPBS 5 times between each step of the assay. Clones were considered positive if the optical density 450 nm was greater that two standard deviations above the background.

### Bio-layer Interferometry Selection of Receptor Binding Domain Binding VHH Clones

ELISA-positive clones were further assessed by bio-layer interferometry binding to RBD. Assay buffer was defined as 0.1% BSA (w/v) in 1xPBS. All assay conditions were prepared in a Greiner 96-well plate (#655209) in a volume of 300 µL. Biotinylated SARS-CoV-2 S protein RBD (SPD-C82E9, ACRO Biosystems) with a C-terminal AviTag was used to ensure uniform directionality of the protein. Biotinylated-RBD was diluted into assay buffer at 1 µg/mL and immobilized onto streptavidin coated biosensors (#18-5019, BioForte) to a minimum response value of 1nm on the Octet Red96 System (ForteBio). A baseline response was established in assay buffer prior to each association. Candidate clone supernatant prepared previously was diluted 1:1 with assay buffer and allowed to associate for 60s and dissociate for 60s, which provided a sufficient interval to detect positive binding. Biosensors were regenerated in 250 mM imidazole in assay buffer for 45s which removed residual bound candidate nanobody and returned to the baseline well. This method allowed for detection of clones that bind RBD in both ELISA and a qualitative estimate of binding based on response curve dynamics. Clones positive on both the RBD ELISA and bio-layer interferometry measurement were selected for sequencing and further characterization.

### ProteOn XPR36 surface plasmon resonance affinity measurement of NIH-CoVnb-112 RBD binding

Surface plasmon resonance affinity and kinetic measurements were performed using the ProteOn XPR36 (Bio-Rad). Lanes of a general layer compact (GLC) chip were individually coated with 2 µg/mL SARS-CoV-2 Receptor Binding Domain (RBD) (#SPD-C52H3, ACRO Biosystems) in 10 mM sodium phosphate pH 6.0 and attached to the chip following the standard 1-ethyl-3-(3-dimethylaminopropyl)carbodiimide hydrochloride (EDC)/*N*-hydroxysulfosuccinimide (sulfo-NHS) coupling chemistry available from the manufacturer resulting ∼1000RU of protein deposited. Binding kinetics of NIH-CoVnb-112 were tested at 25 °C by flowing six concentrations varying from 300 to 0 nM at 100 μL/min for 90 seconds or more then dissociation was monitored for at least 600 seconds. Following each run, the chip was regenerated by flowing 0.85% phosphoric acid (∼ pH 3.0) across the surface. Data analysis was performed with ProteOn Manager 2.1 software, corrected by subtraction of an uncoated column as well as interspot correction. The standard error of the fits was less than 10%. Binding constants were determined using the Langmuir model built into the analysis software.

### Sanger Dideoxy Sequencing

Clones enriched by phage display and positively selected by ELISA assay and bio-layer interferometry measurement were sequenced using a universal Lac-promoter primer (Lac-fwd: 5’-CGTATGTTGTGTGGAATTGTGAGC-3’) with standard Sanger dideoxy sequencing at Genewiz. The resulting sequences with Quality Scores above 40 and with Contiguous Read Lengths of greater than 500 were considered reliable. Sequences were trimmed to include only the VHH coding region and the protein coding sequences aligned using the Clustal Omega algorithm[40] included in SnapGene software (GSL Biotech LLC).

### Nanobody expression and purification

Lead candidates were expressed by transferring the phagemid from TG-1 cells into the BL21 (DE3) competent *E*. *coli* strain (C2527I, New England BioLabs). Cultures were grown in 250mL 4xYT-carbenicillin in a 1-liter baffled flask at 37 °C and 300rpm until the OD600 reached between 0.7-0.9. Protein expression was induced by addition of isopropyl-beta-D-thiogalactoside to a final concentration of 1mM and the culture conditions reduced to 30 °C and 180rpm for overnight growth. The expression cultures were pelleted at 8000xG for 10 minutes at 4 °C; the resulting pellets maintained on ice. Osmotic shock was achieved by resuspension of the cell pellet in TES buffer (0.2M Tris, 65µM EDTA, 0.5M sucrose, pH 8.0) at 1/5^th^ the original culture volume (e.g. 20 mL TES for 100mL culture volume) and incubated on ice for 1 hour with occasional agitation. The suspension was pelleted again as described above and then resuspended in an equal volume of ice-cold 10 mM magnesium chloride and incubated on ice for 30 minutes with occasional agitation. A final centrifugation was performed at 17,000xG for 10 minutes at 4 °C to pellet all cellular debris. The resulting supernatant was filtered through a 0.22 µm syringe filter to remove any residual particulates.

Purification of the nanobody was accomplished by injection of the clarified osmotic shock supernatant onto a 1mL HisTrap FF column (#17531901, Cytiva Lifesciences) attached to an AKTA Pure FPLC system. Unbound protein was washed with 10 column volumes 1xPBS followed by 10 column volumes 20 mM imidazole in 1xPBS. The enriched nanobody was eluted with 250 mM imidazole in 1xPBS. An automated collection of the unbound flow-through and fractionation of the wash and elution steps was achieved with the fraction collector F9-C. The nanobody typically eluted into two 1.5 mL fractions which were pooled and concentrated to a volume of approximately 1mL using a 10 kDa molecular weight cut-off Centricon centrifugal concentrator (#MPE010025, EMD Millipore). The concentrated volume was then injected onto a Superdex 75 10/300 GL size exclusion chromatography column on the AKTA Pure system with 1xPBS as the eluent and fractions collected. This polishing step typically achieved a >90% purity as determined by SDS-PAGE. Protein concentration was estimated using absorbance at 280 nm and quantified using a standard Bicinchoninic acid protein assay (#23225, ThermoScientific) with BSA as the concentration standard.

### Bio-layer Interferometry affinity measurement of RBD binding VHH clones

Bio-layer interferometry was used to measure the affinity binding constants of purified VHH clones. Assay buffer is defined as 0.1% BSA (w/v) in 1xPBS. All assay conditions are prepared in a Greiner 96-well plate (#655209) in a volume of 300 µL. Biotinylated SARS-CoV-2 S protein RBD (SPD-C82E9, ACRO Biosystems) with a C-terminal AviTag was used to ensure uniform directionality of the protein. Biotinylated-RBD was diluted into assay buffer at 1 µg/mL and immobilized onto streptavidin coated biosensors (#18-5019, BioForte) to a minimum response value of 1 nm on the Octet Red96 System (ForteBio). A baseline response was established in assay buffer prior to each association. The purified VHH clones were diluted into assay buffer at the specified concentrations (typically 1,000 nM to 0 nM). The VHH clones were allowed to associate for 180-240s followed by dissociation for 300-600s in the same baseline wells. The assay included one biosensor with only assay buffer which was used as the background normalization control. Using the ForteBio Data Analysis suite, the data was normalized to the association curves following background normalization and Savitzky-Golay filtering. Curve fitting was applied using global fitting of the sensor data and a steady state analysis calculated to determine the association and dissociation constants.

### Receptor Binding Domain Competition Assay

SARS-CoV-2 RBD (#SPD-C52H3, ACRO Biosystems) was coated at 0.5 µg/mL in sodium carbonate buffer (pH 9.6), 100 µL per well, onto Nunc Maxisorp plates overnight at 4°C. Coating solution was removed, and plate blocked with 300 µL 2% BSA (#5217, Tocris) in 1xPBS for 1 hour at room temperature. Purified nanobodies were diluted at specified concentrations in 0.2% BSA in 1xPBS and transferred to the blocked plate in triplicate. The nanobodies were incubated for 45 minutes to allow for association with RBD. Biotinylated ACE2 (AC2-H82E6, ACRO Biosystems) was prepared at 0.2 µg/mL in 0.2% BSA solution. 10 µL of the ACE2-biotin solution was transferred into each well of the assay and allowed to incubate for 15 minutes with at 600 rpm. The assay plate was washed and 100 µL of poly-streptavidin (#85R-200, Fitzgerald Industries International) diluted 1:2000 in 2% BSA solution was transferred to each well and incubated with at 600 rpm for 30 minutes. After a final wash the plate was developed by addition of 100 µL tetramethylbenzidine (#T5569, Sigma-Aldrich). Assay development was stopped by addition of 50 µL 1M sulfuric acid and the assay absorbance measured at 450 nm on a Biotek Synergy 2 plate reader. The assay plate was washed with 1xPBS 5 times between each step of the assay.

### Receptor binding domain variant competition assay

SARS-CoV-2 RBD variants have been noted which increase the affinity for human ACE2 binding. Competition between the RBD mutants was performed as described above with the novel mutation RBD proteins replacing the canonical RBD sequence. SARS-CoV-2 RBD mutants W436R, N354D/D364Y, and V367F (#SPD-S52H7, SPD-S52H3, and SPD-S52H4 respectively, ACRO Biosystems) were coated at 0.5ug/mL in sodium carbonate buffer (pH9.6), 100µL per well, onto Nunc Maxisorp plates overnight at 4°C. The assay was completed as described above.

### Genscript SARS-CoV-2 Neutralization Antibody Detection Kit

Secondary assessment of the RBD-ACE2 interaction blocking potential of the isolated nanobodies was performed using the Genscript SARS-CoV-2 Neutralization Antibody Detection Kit (#L00847, Genscript) according to the manufacturer’s instructions. Briefly, horse radish peroxidase conjugated RBD was diluted using the supplied assay buffer and then incubated with specified concentrations of nanobody for 30 minutes at 37 °C. The samples were then transferred onto the supplied human ACE2-coated assay plate and incubated for 15 minutes at 37°C. The plate was washed using the supplied solution and the assay developed using a supplied 3,3′,5,5′-Tetramethylbenzidine reagent. The assay was stopped with 50 μL of the supplied Stop Solution. The assay was measured at 450 nm on a Biotek Synergy 2 plate reader.

### SARS-CoV-2 pseudotyped lentivirus-based transduction fluorescence inhibition assay

All lentiviruses were propagated in HEK293T/17 cells (ATCC # CRL-11268) according to published Current Protocols in Neuroscience.[41] Briefly, 293T cells were transiently transfected with plasmids expressing SARS-CoV-2 spike protein (GenScript MC_0101081, human codon optimized, ER retention signal removed), psPAX2 (Addgene #12260), and a lentiviral transfer vector CD512-EF1a-RFP (System Biosciences CD512B-1) using Lipofectamine 2000. Supernatant was collected 48 hours post transfection and concentrated by centrifugation at 50,000 g for 2 hours over a 20% sucrose cushion. Pellets were resuspended in PBS and used for infection. All titers were determined by performing biological titration of fluorescent viruses by flow cytometry.

For transduction assays, HEK293Ts expressing human Angiotensin-Converting Enzyme 2 (HEK293T-ACE2, BEI Resources, NR-52511) were plated at the density of 50,000 cells/well in a 6-well plate. Cells were transduced with SARS-CoV-2 pseudotyped recombinant lentiviruses expressing RFP (S-CD512-EF1a-RFP) with multiplicity of infection (MOI) of 0.5 +/-nanobodies at described concentrations. Media on cells was replaced the next day. 48 hours post transduction, cells were released from wells with trypsin and fixed in 1% formaldehyde. BD LSRFortessa™ Flow Cytometer was used to determine percent fluorescent cells and the mean fluorescent intensity per sample. All experiments were performed in triplicate.

### Expression of NIH-CoVnb-112 in *Pichia pastoris*

Increased expression yield and elimination of potential endotoxin from the bioprocess was achieved by cloning and expression of the nanobody in the methylotrophic yeast *Pichia pastoris*. The EasySelect Pichia Expression Kit (#K1740-01, Invitrogen) was used for cloning of the VHH sequence into expression vector pICZα by amplification of the pHEN2 containing phagemid with reformatting primers (pichia-nominal-fwd : 5’-TATCTCTCGAGAAAAGAGATGTGCAGCTGCAGGAGTCTG-3’ and pichia-nominal-rev: 5’-TTGTTCTAGATTAGTGATGGTGATGATGATGTGCGGCCGC-3’). The reformatting primers introduce XhoI and XbaI (New England Biolabs) restriction endonuclease sites which allow for in-frame cloning of the VHH sequence with the α-factor secretion signal and includes a vector independent 6xHis tag at the C-terminus. The resulting expression vector was linearized using ScaI-HF (New England Biolabs) and transformed into competent *Pichia p*. strain X-33 followed by selection on YPD agar with 1M sorbitol and 100µg/mL Zeocin. Resulting clones were confirmed for recombination and for phenotype by small scale expression. For scaled expression, selected clones were grown in buffered glycerol complex medium to an OD600 of 1 prior to addition of methanol to a final concentration of 0.5%. The culture was batch-fed every 24 hours to a final methanol concentration of 0.5% until harvest of the culture supernatant. Spent cells were removed by centrifugation at 3000xG for 10 minutes at room temperature and the supernatant clarified by filtration through a 0.45µm filtration unit. The media supernatant was concentrated using a Minimate™ tangential flow filtration system (#OAPMPUNV, Pall) and buffer exchanged using 1xPBS containing 10mM imidazole. Purification was performed as described above using Ni-NTA affinity chromatography.

### Nebulization stability assessment of NIH-CoVnb-112

Stability following nebulization of *Pichia pastoris* expressed NIH-CoVnb-112 was performed using an Aerogen® Solo High-Performance Vibrating Mesh nebulizer placed in line with a custom glass bead condenser. A plastic culture tube was fitted with a glass-pore frit and filled with sterilized 5mm borosilicate glass beads. A three-way stopcock was positioned distal to the frit to prevent pressurization during nebulization. A 2mg/mL NIH-CoVnb-112 solution was prepared in 0.9% normal saline to model potential patient delivery. The nanobody was nebulized and the resulting condensate incubated at 37°C for 24hr to mimic exposure to body temperature. The nebulized, 37°C treated nanobody was then collected for stability assessments. Pre and post-nebulization samples were denatured in LDS sample buffer (Invitrogen) and run on a NuPAGE 12% Bis-Tris precast polyacrylamide gel with SeeBlue™ Plue 2 protein standards. Additional pre and post-nebulization samples were injected onto a Superdex 75 Increase 10/300 GL size exclusion column operating on an AKTA Pure 25 M system.

### Stability of NIH-CoVnb-112 in human plasma

NIH-CoVnb-112 expressed in *Pichia pastoris* was diluted from a concentrated stock solution into apheresis derived pooled human plasma (#IPLA-N, Innovative Research) to a final concentration of 5µM and incubated at 37°C for either 24hr or 48hr with gentle rotation. An identical sample set was prepared at 5µM in a solution containing 35mg/mL recombinant human albumin (#A9731, Sigma-Aldrich) and incubated at 37°C for either 24hr or 48hr with gentle rotation. A no-incubation control for each the plasma and recombinant human albumin conditions was prepared at 5µM. The samples were prepared in a manner providing all conditions complete at the same time. The samples were diluted 1:10 with 1xPBS to yield a final nanobody concentration of 500nM and Bio-layer Interferometry was performed using immobilized biotinylated SARS-CoV-2 S protein RBD to determine retention of binding potential.

### Determining Melting Temperature and Refolding by Circular Dichroism

Circular Dichroism (CD) was performed using a Jasco J-815 Spectropolarimeter. For thermal stability measurements NIH-CoVnb-112 was diluted to 10 μg/mL in deionized water and placed in a quartz cuvette with 1 cm path length and CD was measured at an ultraviolet wavelength of 205 nm. NIH-CoVnb-112 heated from 25°C to 85°C at a rate of 2.5°C/min while stirring and then cooled back to 25°C at the same rate.

## ACKNOWLEDGEMENTS

We are grateful to Chris Broder, Pat Casey, Dan Chertow, Ed Engleman, Cory Hallum, Alura Johnston, Walter Koroshetz, Steve Jacobson, Avi Nath, and all of the members of the Brody lab for helpful discussions. We thank NHLBI Biophysics Core Facility for the use of BLI instrument. We thank the NIH SARS-CoV-2 committee for approval and support. We thank Jay, Jason, and Donald Jorgensen at Triple J Farms for expedited llama vaccination services. We thank Jerrel Yakel, Neurobiology Lab Chief, for sharing the Viral Vector Core resources for this project. The study was supported by the NINDS intramural research program under the Laboratory of Functional and Molecular Imaging, directed by Alan Koretsky. A provisional patent has been filed (U.S. Provisional Application No.: 63/055,865, Filing Date July 23, 2020).

## AUTHOR CONTRIBUTIONS STATEMENT

DLB and TJE conceived the experiments. TJE, NPM, GPA, and ERG performed experiments. DLB and TJE wrote the manuscript.

---

**Supplementary Fig. 1:**
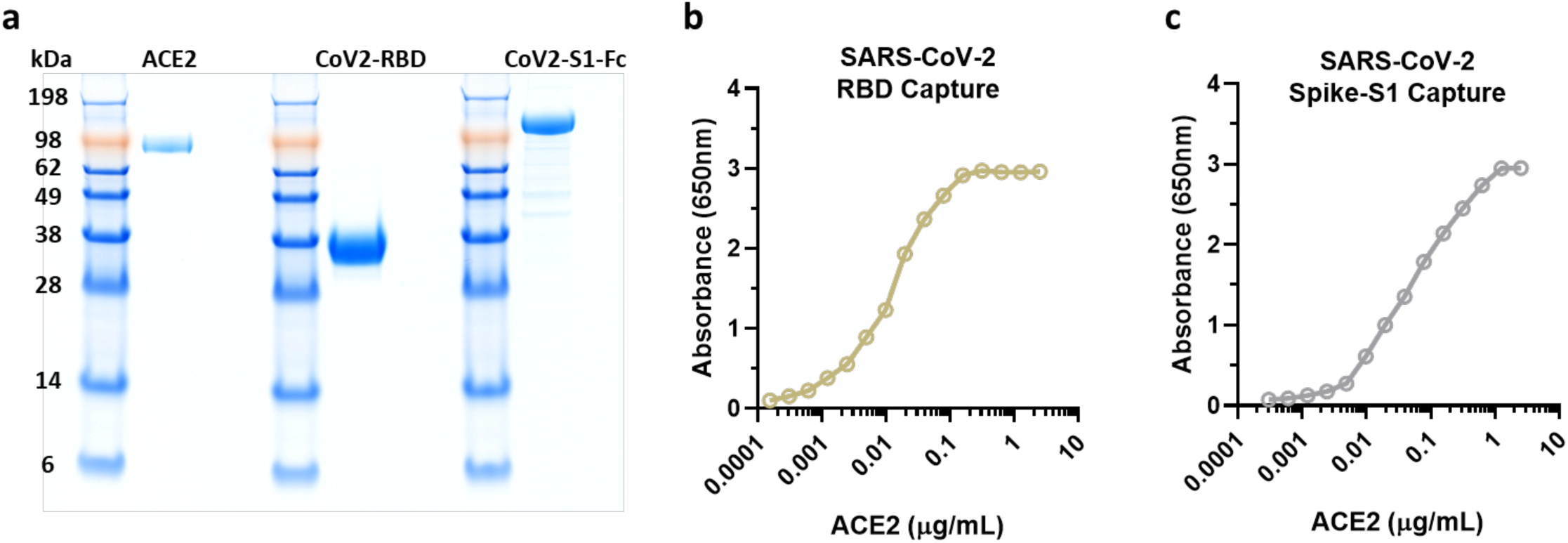
Reagents required for characterization and validation include recombinant human ACE2, recombinant SARS-CoV-2 receptor binding domain (RBD), and recombinant SARS-CoV-2 Spike protein (S1). **a** The presence of prominent, single bands on a SDS-PAGE gel for each protein indicate purity and appropriate size. **b**,**c** Validation that recombinant SARS-CoV-2 RBD and SARS-CoV-2 Spike S1 bind with high affinity to recombinant human ACE2. This high affinity, saturable binding indicates that all 3 recombinant proteins are appropriately folded *in vitro*.

**Supplementary Fig. 2:**
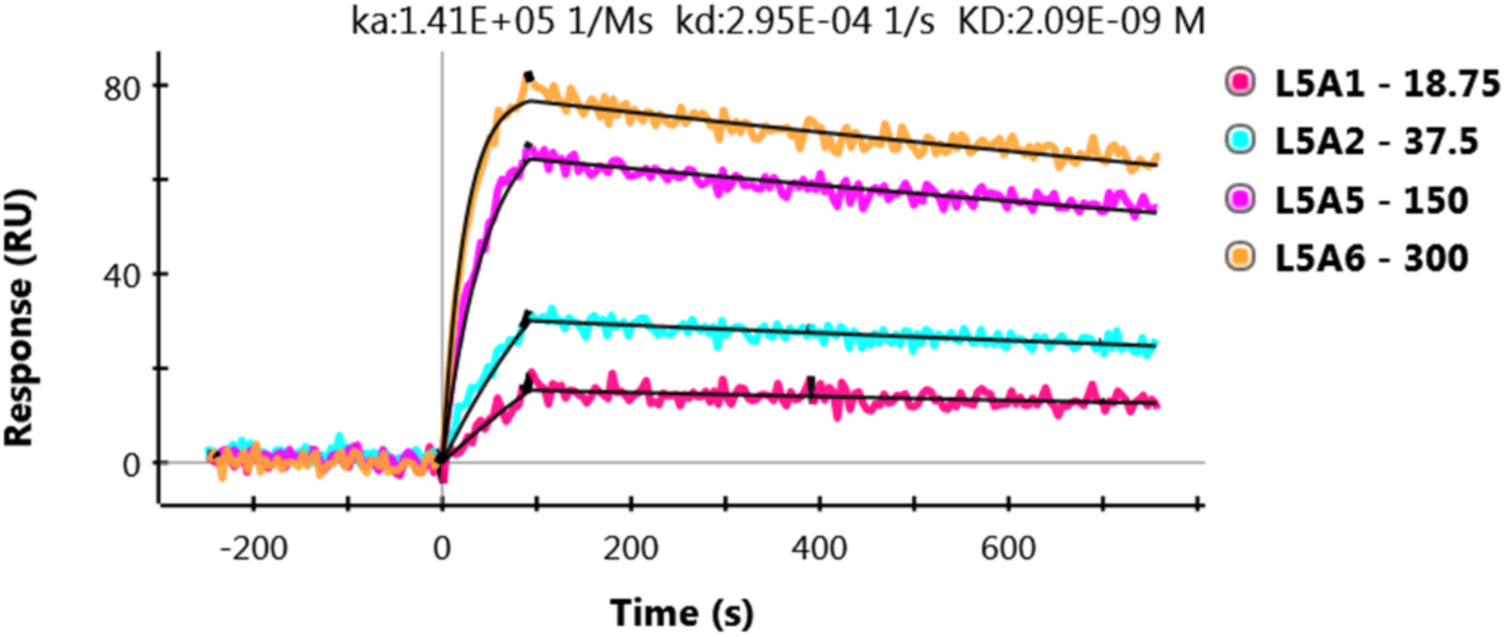
Surface plasmon resonance affinity measurement of NIH-CoVnb-112 binding RBD. Validation of the high affinity binding constant of NIH-CoVnb-112 was established by measurement of binding using an alternative affinity measure. SARS-CoV-2 RBD was immobilized onto the surface of a GLC chip using EDC/NHS chemistry followed by flow of NIH-CoVnb-112 across the chip surface to allow for measurement of binding kinetics. Raw sensor data is overlaid by software curve fits which allow for measurement of the association and dissociation constants using the Langmuir model. The calculated affinity binding constant (2.1nM) is in close agreement with the value measured with biolayer interferometry (4.9nM).

**Supplementary Fig. 3:**
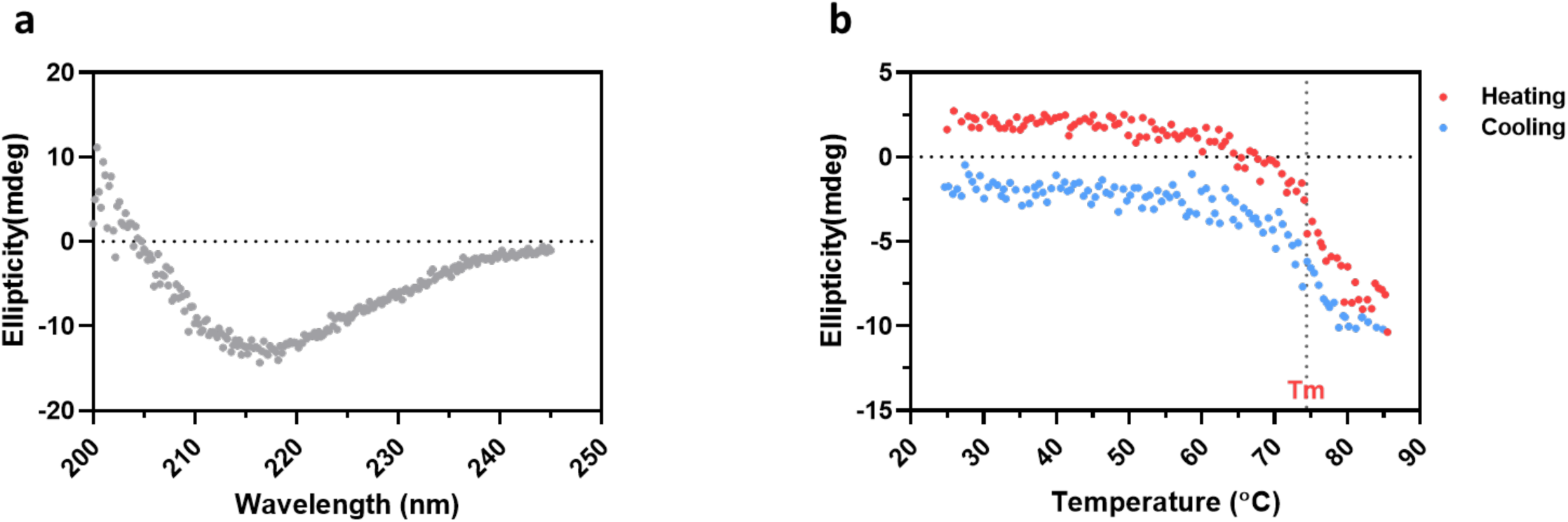
Characterization of NIH-CoVnb-112 by circular dichroism. **a** Representative CD curve for NIH-CoVnb-112. **b** Reversible folding was monitored using circular dichroism using a Jasco J-815 spectropolarimeter at 205nm during a heating-cooling cycle over 25°C to 85°C at a rate of 2.5 °C/min. The inflection point at 74.4°C indicates the melting temperature. Using the delta between the heating and cooling curves is used to calculate a 73% refolding rate for NIH-CoVnb-112 which is indicative of a highly stable structure.

**Supplementary Fig. 4:**
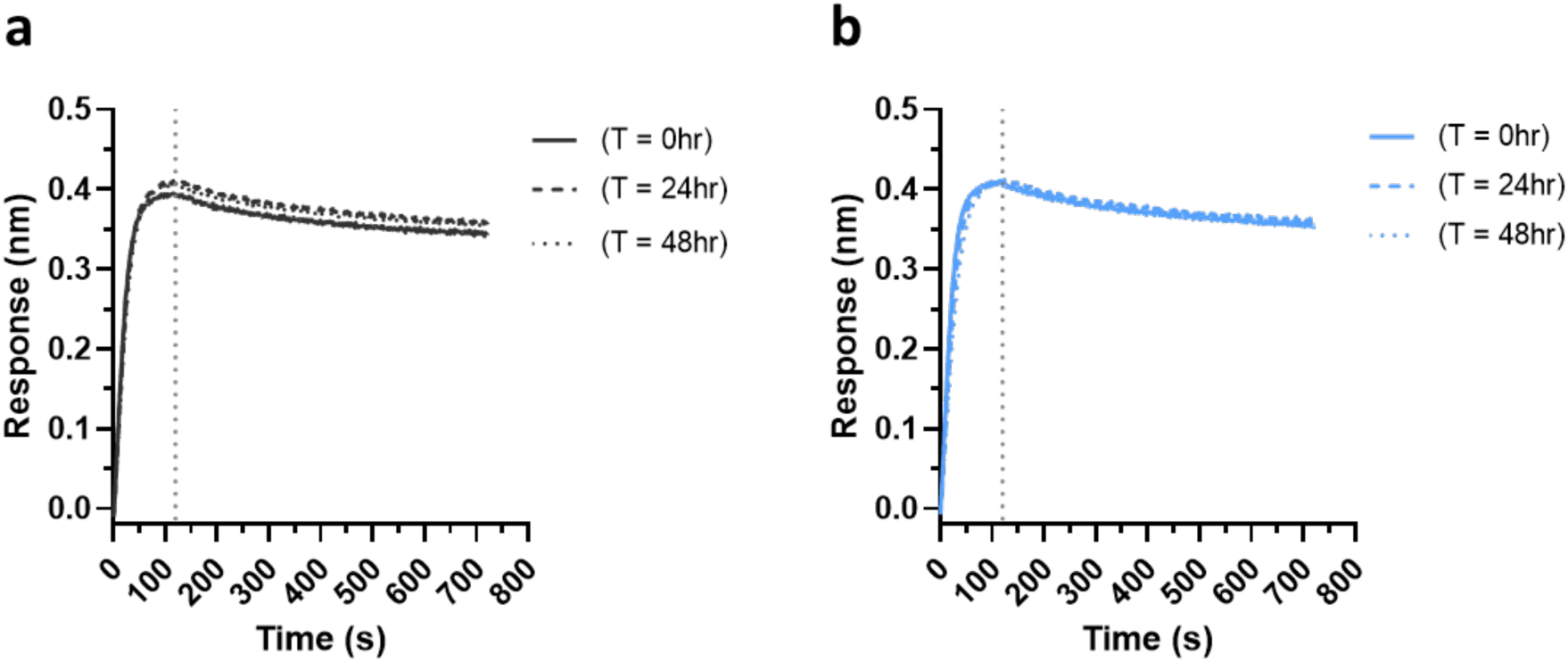
NIH-CoVnb-112 is highly stable during controlled incubation with human plasma and albumin. To determine the if NIH-CoVnb-112 is sensitive to plasma conditions a controlled incubation was performed at 37°C with mixing. Measurement of binding, following incubation for 24 or 48hrs, of NIH-CoVnb-112 in (**a)** pooled human plasma or (**b)** recombinant human albumin alone to mimic conditions which could lead to degradation and loss of function. NIH-CoVnb-112 was additionally spiked into pooled human plasma or recombinant human albumin immediately prior to measurement by biolayer interferometry.

**Supplementary Fig. 5:**
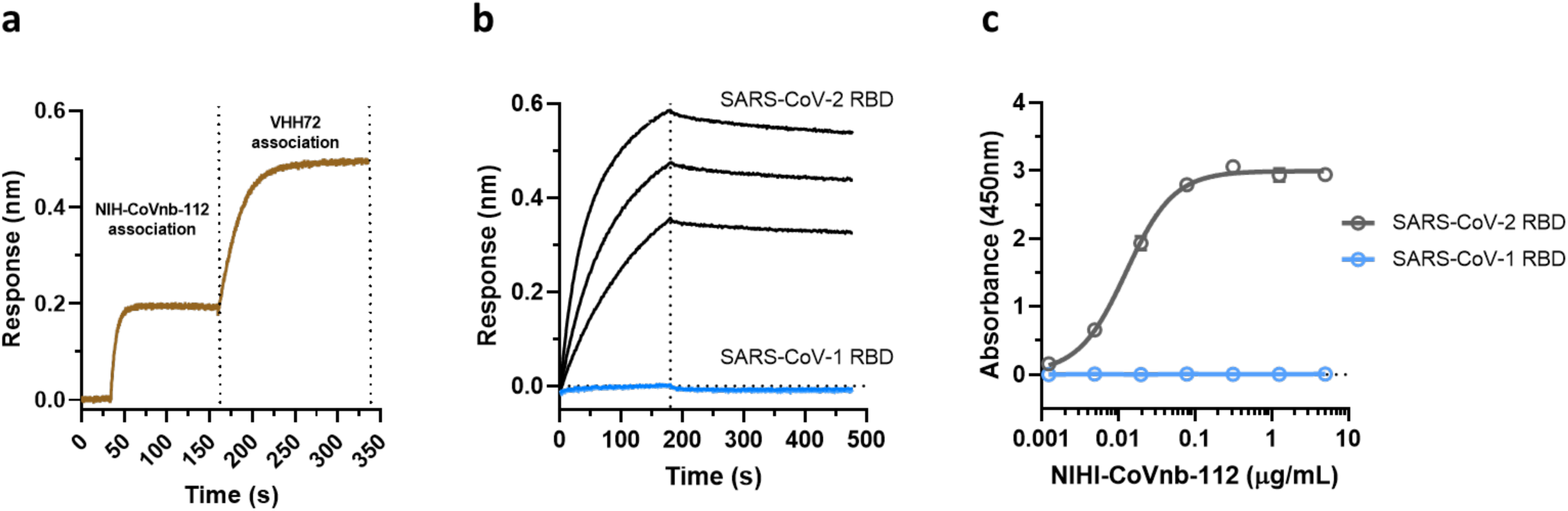
NIH-CoVnb-112 binds to SARS-CoV-2 RBD at a distinct epitope from that bound by VHH72 and does not bind to SARS-CoV-1 RBD. **a** To determine if NIH-CoVnb-112 and VHH72 have competing epitopes the following association Octet experiment was performed: Biotinylated SARS-CoV-2 RBD was bound to streptavidin BLI sensors in blocking buffer. Following baseline stability, the sensor was transferred into a well containing 500nM NIH-CoVnb-112 and allowed to associate. The sensor was then transferred to the adjacent well containing 500nM VHH72 and allowed to associate. If VHH72 had an overlapping epitope blocked by NIH-CoVnb-112 there would be minimal to no increase in response. Based on the observed increase in response, it can be inferred that the two nanobodies have non-competing epitopes on the SARS-CoV-2 RBD. **b** An Octet experiment was performed to determine if NIH-CoVnb-112 possesses permissive binding to SARS-CoV-1 RBD. NIH-CoVnb-112 was biotinylated using NHS-LC-biotin and then immobilized onto a streptavidin biosensor followed by association of SARS-CoV-1 RBD (*blue curves*) and SARS-CoV-2 (*black curves*) at 500, 250, and 125nM RBD. NIH-CoVnb-112 does not bind to SARS-CoV-1 RBD. **c** As an orthogonal confirmation of the Octet experiment, binding was measured by ELISA. SARS-CoV-1 RBD (*blue open circles*) and SARS-CoV-2 RBD (*black open circles*) were coated on to an ELISA plate at 10 micrograms/mL and incubated with a range of NIH-CoVnb-112 concentrations. An anti-alpaca secondary antibody was used for detection and confirms the lack of NIH-CoVnb-112 binding to SARS-CoV-1 RBD.

**Supplementary Fig. 6:**
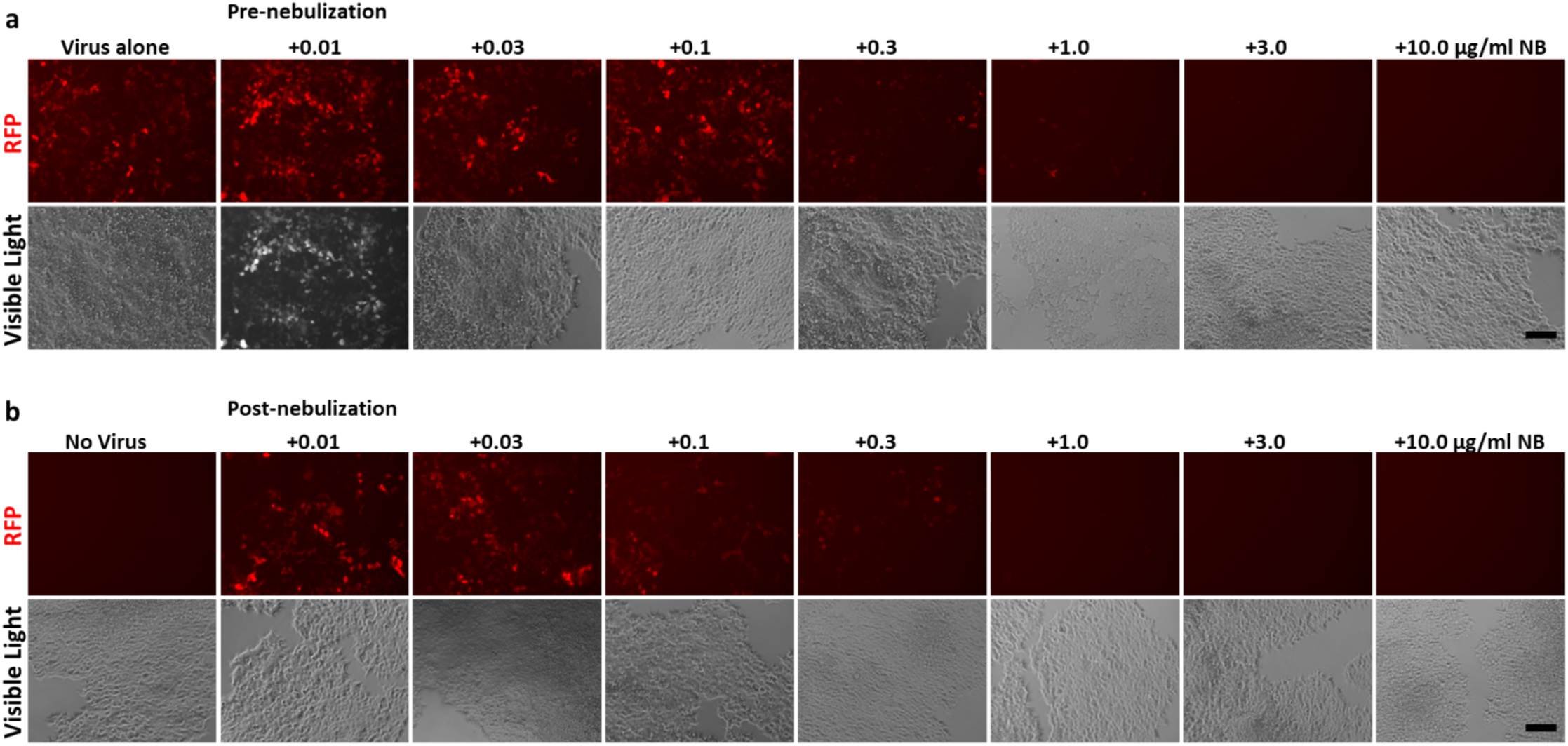
Epifluorescence microscopy of inhibition of SARS-CoV-2 pseudotyped virus transduction by NIH-CoVnb-112. Inhibition of a SARS-CoV-2 pseudovirus transduction was performed (MOI=0.5) on HEK293-ACE2 cells with a range of NIH-CoVnb-112 concentrations from nanobody before and after nebulization to mimic therapeutic inhaled administration. Prior to flow cytometry evaluation, fields in each well were imaged by brightfield and RFP epifluorescence for the (**a**) pre-nebulization and (**b**) post-nebulization samples. At and above concentrations of 1 microgram/mL NIH-CoVnb-112 there is a robust inhibition of fluorescence signal, while below 0.3 micrograms/mL there is a gradual increase to comparable levels to the virus alone control. This demonstrates the potent inhibition of the biologically active SARS-CoV-2 spike protein and the interaction with its receptor. (Scale bar = 100µm)

**Supplementary Fig. 7:**
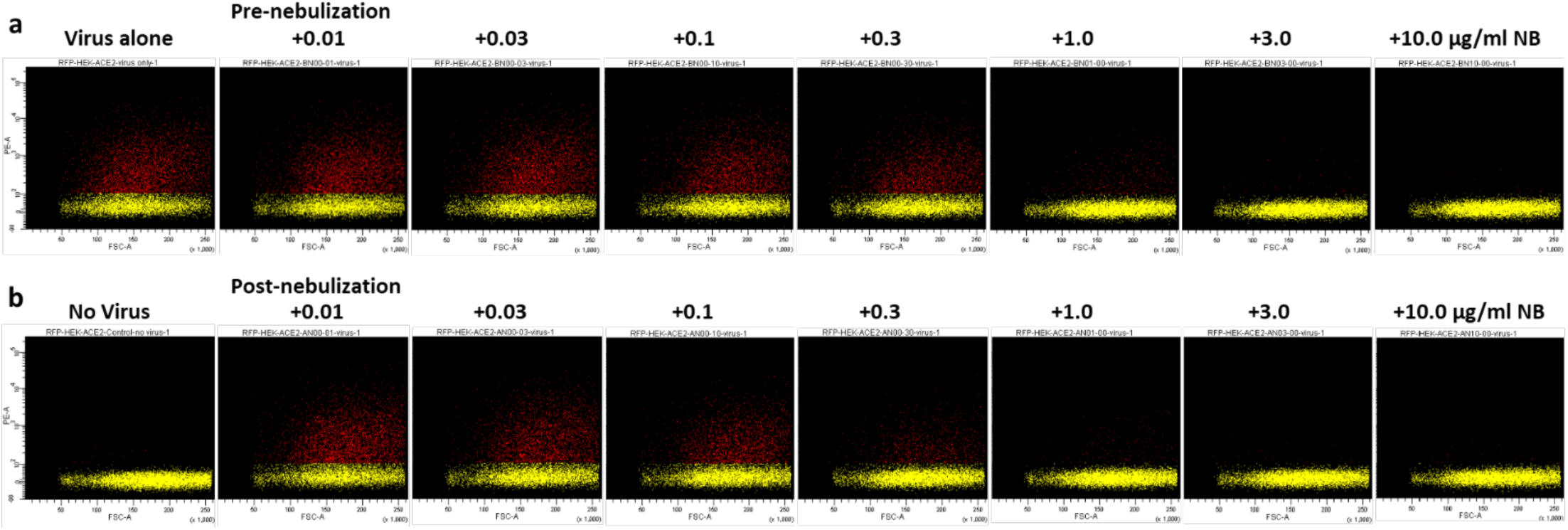
Flow cytometry analysis of HEK293-ACE2 cells following inhibition of SARS-CoV-2 pseudotype virus by NIH-CoVnb-112. Following a 48hr transduction of SARS-CoV-2 pseudovirus in the presence of pre and post-nebulization NIH-CoVnb-112, at various concentrations, the cells were trypsinized and fixed prior to flow cytometry. A single cell population was gated, to exclude debris, and 10,000 events collected per sample on a BDFortessa in the PE-Cy5.5 channel in triplicate. The (**a**) pre-nebulization and (**b**) post-nebulization sample display a very similar distribution of RFP positive cells at each respective concentration of NIH-CoVnb-112. Yellow dots represent gated non-positive events; red dots represent gated positive events.

